# Evaluation and Design of Genome-wide CRISPR/Cas9 Knockout Screens

**DOI:** 10.1101/117341

**Authors:** Traver Hart, Amy Tong, Katie Chan, Jolanda Van Leeuwen, Ashwin Seetharaman, Michael Aregger, Megha Chandrashekhar, Nicole Hustedt, Sahil Seth, Avery Noonan, Andrea Habsid, Olga Sizova, Lyudmila Nedyalkova, Ryan Climie, Keith Lawson, Maria Augusta Sartori, Sabriyeh Alibeh, David Tieu, Sanna Masud, Patricia Mero, Alexander Weiss, Kevin R. Brown, Matej Usaj, Maximilian Billmann, Mahfuzur Rahman, Michael Constanzo, Chad L. Myers, Brenda J. Andrews, Charles Boone, Daniel Durocher, Jason Moffat

## Abstract

The adaptation of CRISPR/Cas9 technology to mammalian cell lines is transforming the study of human functional genomics. Pooled libraries of CRISPR guide RNAs (gRNAs), targeting human protein-coding genes and encoded in viral vectors, have been used to systematically create gene knockouts in a variety of human cancer and immortalized cell lines, in an effort to identify whether these knockouts cause cellular fitness defects. Previous work has shown that CRISPR screens are more sensitive and specific than pooled library shRNA screens in similar assays, but currently there exists significant variability across CRISPR library designs and experimental protocols. In this study, we re-analyze 17 genome-scale knockout screens in human cell lines from three research groups using three different genome-scale gRNA libraries, using the Bayesian Analysis of Gene Essentiality (BAGEL) algorithm to identify essential genes, to refine and expand our previously defined set of human core essential genes, from 360 to 684 genes. We use this expanded set of reference Core Essential Genes (CEG2), plus empirical data from six CRISPR knockout screens, to guide the design of a sequence-optimized gRNA library, the Toronto KnockOut version 3.0 (TKOv3) library. We demonstrate the high effectiveness of the library relative to reference sets of essential and nonessential genes as well as other screens using similar approaches. The optimized TKOv3 library, combined with the CEG2 reference set, provide an efficient, highly optimized platform for performing and assessing gene knockout screens in human cell lines.

## Introduction

The generation of gene knockouts is a cornerstone of functional genomics research. The application of CRISPR technology to induce targeted DNA double-strand breaks in mammalian cells [1] coupled to the ability of the endogenous cellular DNA repair machinery to introduce indels when repairing these lesions, has led to the rapid development of pooled-library CRISPR knockout screens in mammalian cells for functional genomics, chemogenomics, the identification of cancer cell vulnerabilities, and other applications [2-10].

CRISPR screens represent a major advance over pooled-library shRNA screens [11], the prior state-of-the-art in mammalian functional screening approaches, in both sensitivity and specificity. The current designs of large-scale CRISPR experiments benefited from the many lessons learned in shRNA screening. In particular, the design of early CRISPR libraries to include several gRNAs targeting each gene has been driven by experience with pooled library shRNA screens [12,13], as well as the unknowns surrounding the application of CRISPR technology in human cells on a large scale. Integrated analysis of multiple reagents targeting the same gene should overcome the noise introduced by variable reagent effectiveness and the unknown frequency and impact of off-target effects.

With several panels of whole-genome cell-line screens published [2, 5, 8-10], we have the opportunity to undertake a meta-analysis as a means to uncover the drivers of screen quality and variability. We re-analyzed sets of CRISPR screens conducted in adherent and suspension cell lines using three different large-scale libraries and evaluated each for quality and consistency. Based on these observations, we refined our list of core essential genes - i.e. the set genes that are likely to be essential in virtually all cell lines. We evaluated the impact of experimental design, including library size, number of replicates, and use of non-targeting controls, on screen performance. Finally, we derived a sequence signature for highly effective guide RNAs and designed an optimized, genome-scale CRISPR library for efficient screening of human cell lines.

## Results

### An updated set of gold-standard essential genes

We applied our BAGEL analysis pipeline [14] to panels of pooled library CRISPR dropout screens from three different groups, each of which used their own custom library [2, 3, 5, 8] (Supplementary Table 1). Additional genome-scale pooled screens were recently published, but those were not included in this initial analysis [9, 10]. Using the previously published reference sets of essential and nonessential genes [15], we classified essential and nonessential genes from seven adherent cell lines screened with the TKOv1 library [2, 16], four suspension cell lines using the Sabatini library [5], and five suspension plus one adherent line screened with the Yusa library [3, 8], for a total of 17 whole-genome screens (Figure 1a and Supplementary Table 1). Though the screens were performed with different libraries and carried out in different labs, the experimental designs are largely similar, with each screen involving a pooled lentiviral library infection of a large number of cells, serial passaging over two to three weeks, PCR amplification of gRNA integration events, and comparison of the relative abundance of endpoint gRNAs to those of a control timepoint collected shortly after infection.

**Figure 1.**
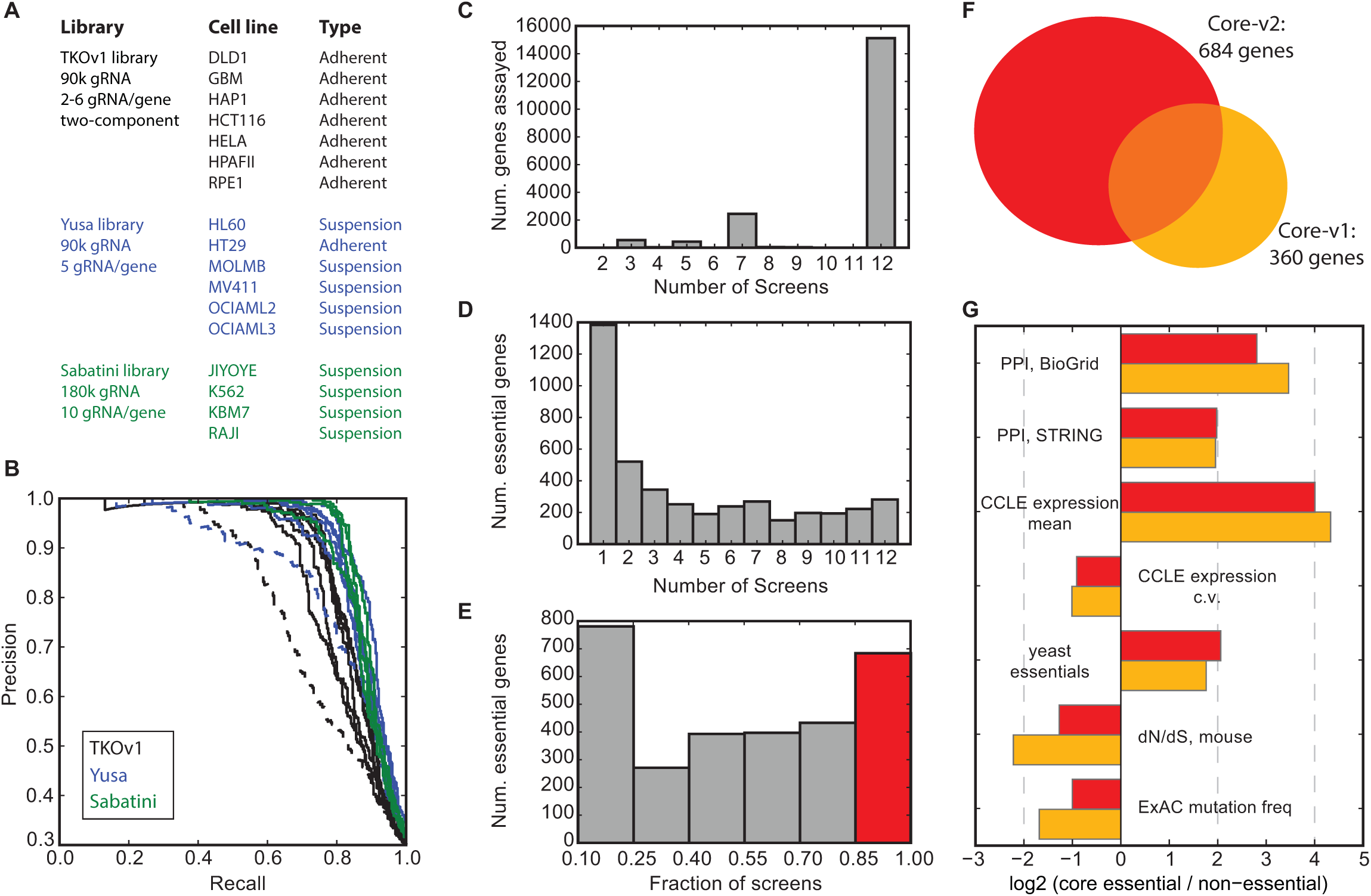
(A) List of CRISPR knockout screens used for this study. (B) Precision-recall curves for the screens in (A) using gold standards defined in [15]. Dashed lines represent low-performing screens that were excluded from further analysis. (C) Number of genes assayed by at least 3 gRNA/gene, across the 12 screens. (D) Number of genes classified as essential (BF>6, FDR<3%) across the 12 screens. (E) Fraction of screens in which a gene is classified as essential. Genes assayed in 7+ screens and essential in 85% of screens (red) are core essentials v2 or CEG2. (F) CEG2 (n=684) is substantially larger and only overlaps CEG1 (n=360; [15]) by ~50%. (G) Functional characterization of CEG2 (Core-v2) vs. CEG1 (Core-v1).

We removed two screens for relatively poor performance (HeLa, MV411) and withheld an additional three for validation studies (HT29, RPE1, K562)(Figure 1b). With the remaining 12 high-performing screens (including 5 adherent and 7 suspension cell lines), we defined an updated set of core essential genes. We defined a gene as being effectively assayed in a cell line if it was targeted by at least 3 independent gRNAs; most genes were assayed in all 12 screens, based on library representation (Figure 1c). Genes that were assayed in at least 7 cell lines and were classified as essential (Bayes Factor, BF, >=6 at FDR <= 3%; Figure 1d) in all, or all but one, of them (max 1 putative false negative allowed) were defined as Core Essential Genes-2.0 or CEG2 (Figure 1e and Supplementary Table 2).

Compared with our previously-defined Core Essentials-1.0 or CEG1, which were derived from a panel of pooled-library shRNA screens [15], CEG2 is 90% larger (n=684 for v2 vs. n=360 for v1; Figure 1f), due largely to the increased sensitivity of CRISPR screens in identifying essential genes at moderate expression levels [2]. However, the CRISPR-derived CEG2 set only include half of the shRNA-derived CEG1 set (183 of 360). We expect that some of the shRNA-specific hits are true essential genes; for example, more than 20 genes coding for ribosomal subunits are included in this list. These genes are not well assayed in most CRISPR libraries due to the difficulty in identifying unique gRNA sequences with low probability of off-target cleavage. However, 131 of the 177 shRNA-only genes are assayed in all 12 CRISPR screens; of these, 26 (20%) were never classified as essential in any CRISPR screen and an additional 24 (18%) were scored as essential in between one and three CRISPR screens. This suggests that the shRNA-only genes may contain a significant number of false positives, possibly resulting from off-target effects on other essential genes. On the other hand, the 501 CRISPR-only additions to the CEG2 set are highly conserved, constitutively expressed, and are central in protein-protein interaction networks (Figure 1g).

With a strict set of genes included in CEG2, we explored how varying experimental design affected the sensitivity of genome-scale CRISPR screens. We determined the minimum number of gRNAs per gene necessary for a high-quality screen, using the CEG2 as positive controls, after recalculating Bayes Factors for all screens using the new training set (Supplementary Table 3). Using data from the Sabatini screens, which used a library of 10 gRNAs per target gene, we randomly selected subsets of two to seven guide RNAs per gene and re-ran the screens *in silico.* At a fixed BF threshold, the fraction of core essentials (Fig 2a) and total number of hits (Fig 2b) increased continuously as the number of gRNAs per gene was increased, although at a diminishing rate. Most of the increased performance was gained with four gRNAs per gene; additional gRNAs per gene added less than 5% more hits, per gRNA added, to the screens (Fig 2c).

**Figure 2.**
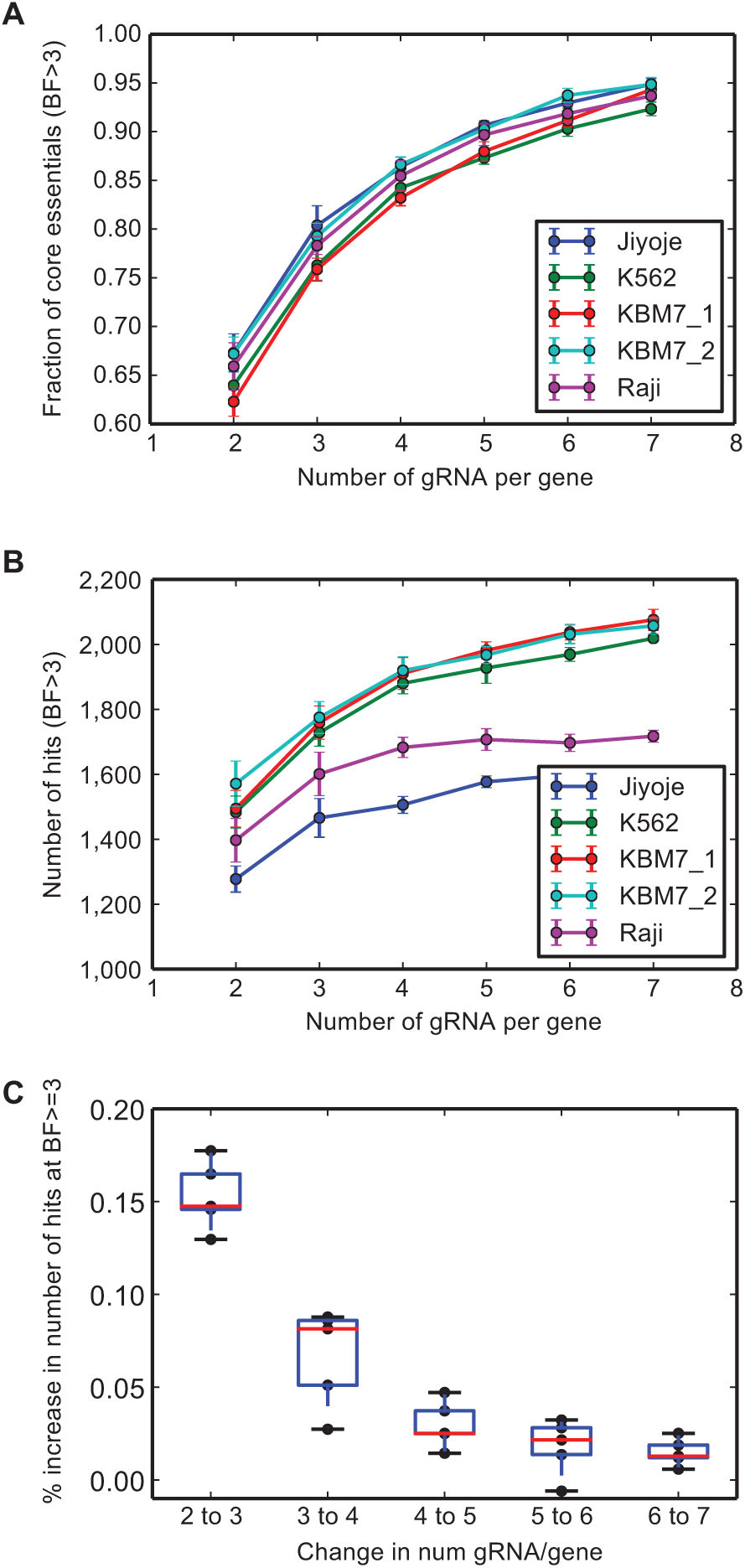
Effect of number of gRNA per gene. (A) Subsets of the Sabatini library were randomly selected and evaluated using BAGEL. The fraction of CEG2 detected is plotted as a function of the number of gRNA/gene. Error bars represent s.d. of 10 random samples from the Sabatini library. (B) Total number of essentials vs. number of guides per gene, from the same samples as (A). (C) Incremental increase in the number of hits vs. incremental increase in the number of gRNA/gene; data from (B).

We also considered the number of replicates for each experiment. Using TKOv1 data from a screen of HAP1 cells and Yusa data from HT29 colorectal cancer cells, which were each performed with three replicates, we measured the performance of one, two (all combinations), or three replicates on screen performance. As expected, additional replicates consistently increased the fraction of core essentials (Fig 3a) and the total number of genes (Fig 3b) called as hits in each screen, but again with diminishing returns: the second replicate increased the number of hits by 9-14%, while the third added <5% (Fig 3c).

**Figure 3.**
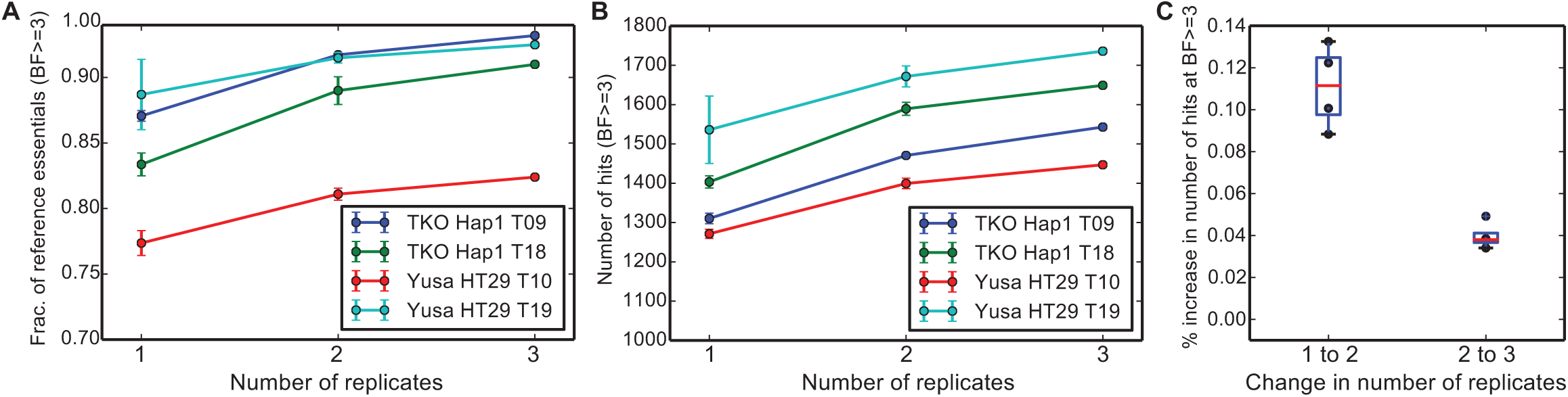
Effect of number of replicates per experiment. The TKOv1 screen in HAP1 cells and the Yusa screen in HT29 cells, each screened at multiple timepoints, were re-analyzed using all combinations of one, two, or three replicates per screen.(A) The fraction of Core-2 reference essentials identified vs. the number of
replicates. (B) The total number of hits vs. the number of replicates. (C) The
incremental increase in total hits, from (B), as the number of replicates is increased.

Finally, we examined the use of nontargeting controls vs. controls that target known or suspected nonessential genes. We identified nontargeting 1,014 control guides in the Sabatini library, whereas we have previously defined a set of nonessential genes for use with training BAGEL and evaluating screen quality [15]. We explored whether there were differences between these two sets of guides. To our surprise, non-targeting controls showed significantly different fold-change distributions than those of guides targeting nonessential genes (Figure 4a-d). Since fold-change is calculated by normalizing read counts and then comparing frequencies, the largest population of minor-phenotype gRNAs will have calculated fold-changes ~0. With approximately eight-fold more gRNAs targeting nonessential genes than nontargeting controls, the larger population has a fold-change distribution centered around zero, while the smaller population appears to have a positive fold-change. In truth, the nontargeting controls likely reflect wildtype growth, while the nonessential controls reflect some small fitness defect from Cas9-induced cleavage and DSB repair, without any locus-specific phenotype. Guides targeting genes with a knockout fitness phenotype will have some combination of locus-specific and nonspecific fitness defect. Thus, when measuring gene-specific fitness effects, it appears that gRNAs inducing DNA double-strand breaks without a gene-specific growth phenotype are a more robust negative control set than nontargeting gRNAs.

**Figure 4.**
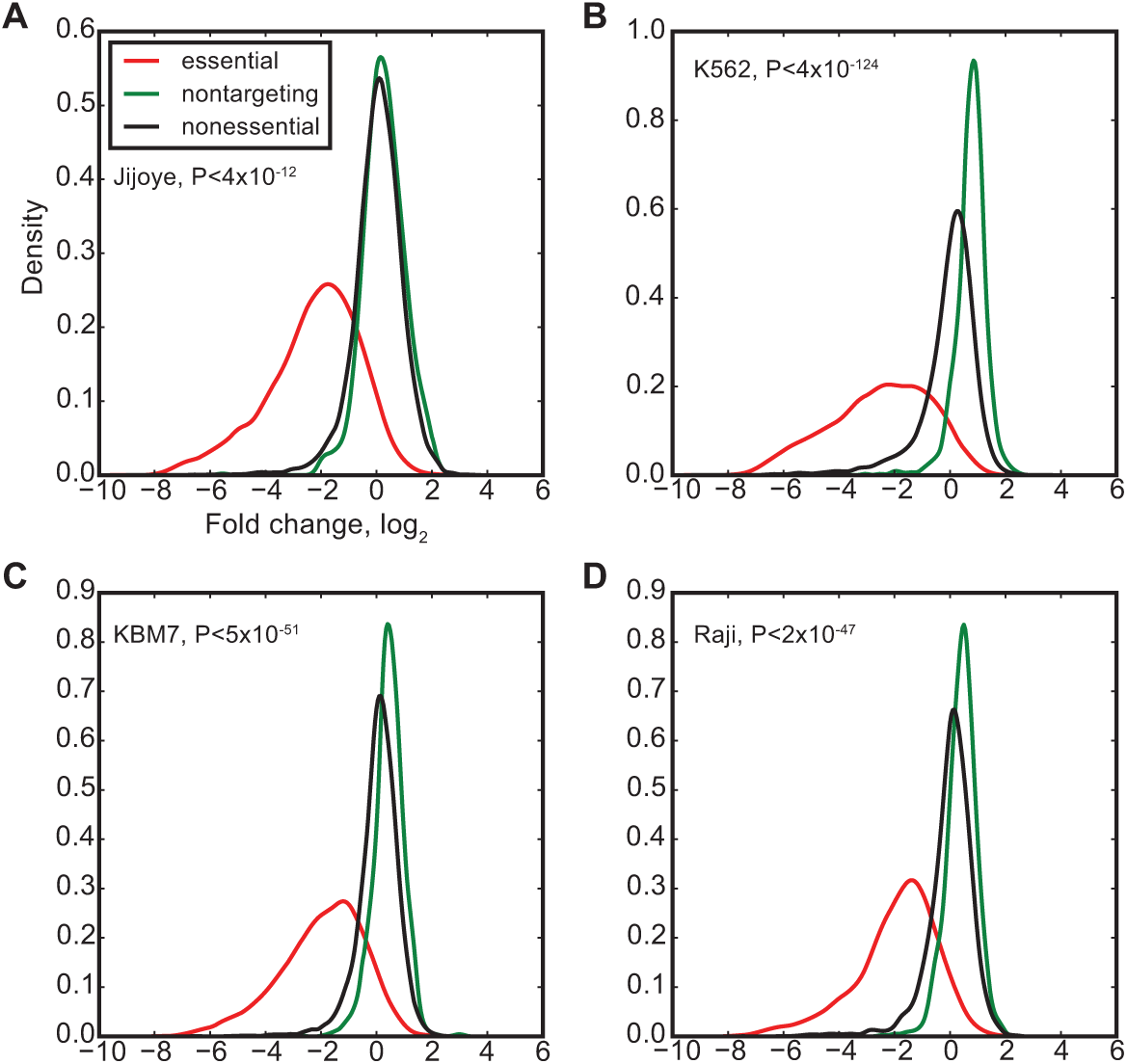
Nonessentials vs. nontargeting controls. The distribution of observed fold-changes of gRNA targeting nonessential genes (black) is compared to the distribution for nontargeting control gRNA (green), in the Sabatini screens of Jiyoye (A), K562 (B), KBM7(C), and Raji (D) cells; P-value from T-test. For reference, the fold-change of gRNA targeting essential genes is also shown (red).

### Designing an optimized library

Given a set of expected outcomes – all core essential genes should drop out of a population in a pooled library screen – we sought to design a sequence-optimized gRNA library that takes advantage of the experimental design characteristics outlined above. Though a variety of small and medium scale experiments have been used to guide gRNA selection algorithms [17-23], our design is, to our knowledge, based on the largest available set of empirical screening results. Using endpoint data from six TKOv1 screens (DLD1, GBM, HAP1, HCT116, RPE1, plus RPE1dTP53, an RPE1-derived cell line), we selected core essential genes targeted by six gRNAs that are each represented by at least 30 reads in the T0 control sample (n=263-360 genes). We rank-ordered the gRNAs for each gene by fold-change and separated the top 3 (“best”) and bottom 3 (“worst”) into separate lists (numbering 789-1,077 gRNA each). We calculated the nucleotide frequency at each position among the best and worst guides across all the screens, subtracted the worst from the best, and normalized the table such that the maximum score at a nucleotide position =1 (see Methods). As expected, the most influential position for gRNA sequence activity is a strong bias toward C at position 18 (Figure 5a and Supplementary Table 4).

**Figure 5.**
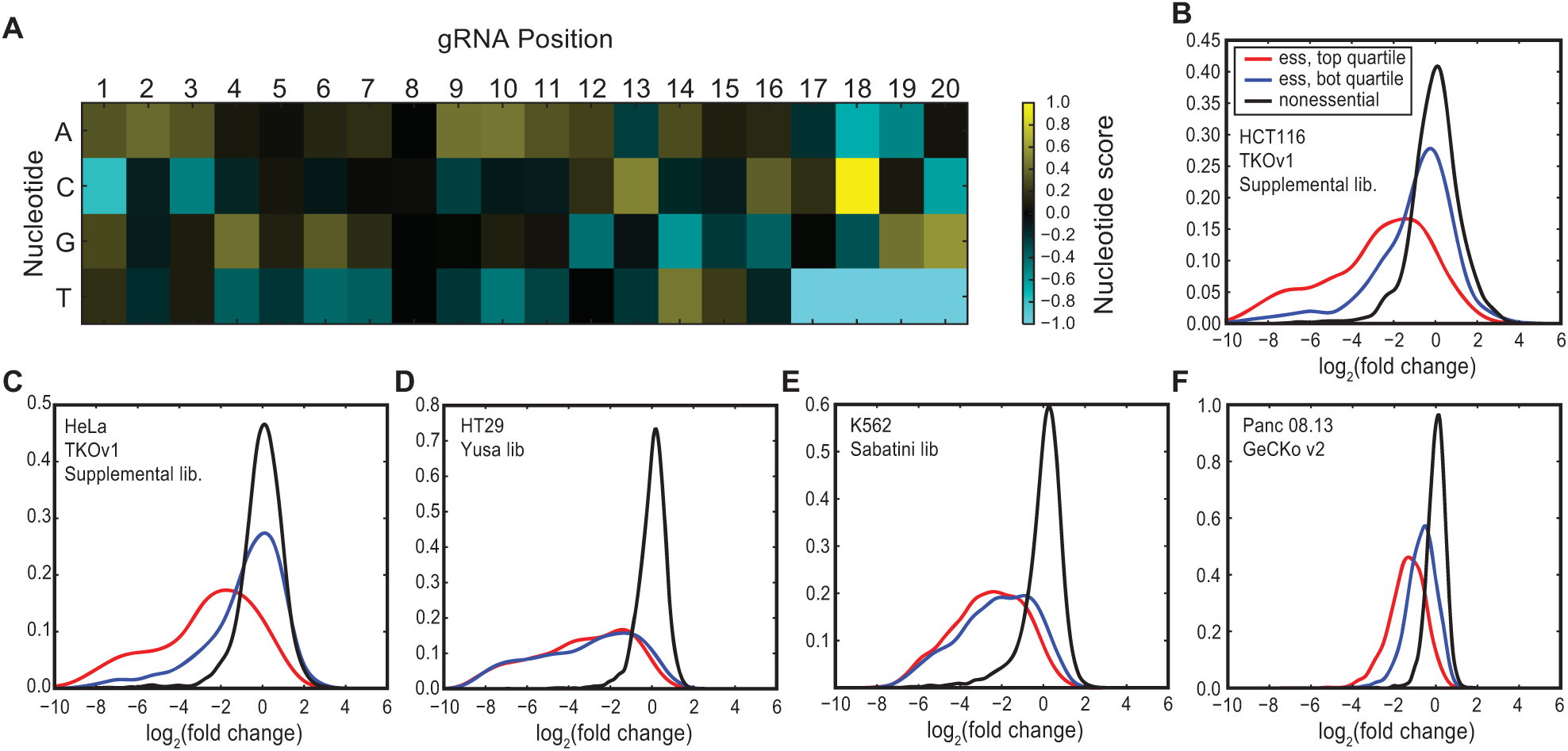
Sequence signature of high-performing guides. (A) Heatmap of the guide score derived from high-performing guides in TKOv1 screens. (B) Across the TKOv1 supplemental library in the HCT116 screen, gRNA targeting CEG2 with sequence scores in the top quartile (red) are compared with gRNA with scores in the bottom quartile (blue). Black, guides targeting nonessential genes. (C-F) Similar plots for TKOv1, Yusa, Sabatini, GeCKOv2 libraries.

To validate the scoring scheme, we used data from TKOv1 screens that included the 85k supplemental library (HCT116 and HeLa cell lines), which added an additional six gRNAs/gene for most genes [2]. We assigned a total score to each gRNA based on the sum of the scores at each nucleotide position in the table, where positive scores indicate a better match to the ‘ideal’ sequence in the score table and negative scores indicate a worse match. We then took the subset of guides targeting core essentials and looked at the fold-change distribution in the top and bottom quartiles of scores within that subset. High scores clearly predicted better performing guides (P-value=1.04×l0^−38^ (HCT116) and 4.83×l0^−50^ (HeLa), T-test; Figure 5b,c) while having minimal difference on nonessential genes in both samples (data not shown). Interestingly, no sequence bias is observed in the Yusa screens (Figure 5d; P-value=0.12). The Yusa library design already includes a sequence optimization step for the last 5 bases (positions 16-20, proximal to the Cas9 PAM sequence) as well as a modified tracrRNA scaffold; these modifications were previously shown to eliminate sequence bias [8] and our analysis is consistent with these results. However, the score table does predict somewhat improved guide performance in the Sabatini screens (Figure 5e; P-value 2.0×10 ^−10^; [5]) and substantially better guides in the Achilles screens (Fig 5f; P-value 1.35×l0^−51^;[10]). Notably, the median gRNA score in the Sabatini library is strongly positive, implying the use of similar design rules, but the median gRNA score in GeCKOv2 is strongly negative. This observation is generally consistent with the substantially better overall performance observed in the Sabatini screens relative to the GeCKOv2 screens, as well as the increased predictive power of our gRNA sequence score for the GeCKO library.

We used the score table to design a sequence optimized CRISPR/Cas9 library that would enable efficient screening of cell lines (see Methods). The Toronto KnockOut version 3 library (TKOv3) is a one-component library (that is, contains Cas9 expression cassette on the plasmid) of 71,090 gRNA with four gRNA/gene targeting 18,053 protein coding genes. The median sequence score of the gRNA in TKOv3 is 1.79, and 97.5% of gRNA have a positive sequence score. In addition, we included 142 gRNA sequences targeting EGFP, LacZ, and luciferase for use as controls in experiments using these reporter genes.

We used the library to screen HAP1 cells and compared the results to TKOv1 screen performed under the same experimental conditions: a single large-scale infection divided into three replicates, with genomic DNA collected after six serial passages. We used the CEG2 set and previously defined reference nonessentials [15] to train and test the BAGEL pipeline for both screens. As shown in Figure 6a, the smaller, sequence-optimized TKOv3 library outperformed the TKOv1 library by precision-recall analysis. It also recovered more essential genes at a strict threshold (BF >6 and FDR < 3%; n=1,850 for TKOv3 vs. n=1,612 for TKOv1), and more of these hits intersect with a list of fitness genes identified in the same cell line using a comprehensive gene trap screen [24] (n=1,534 for TKOv3 vs. n=1,255 for TKOv1, out of n=2,352 fitness genes at 5% FDR identified in [24] Blomen et al; Figure 6b).

**Figure 6.**
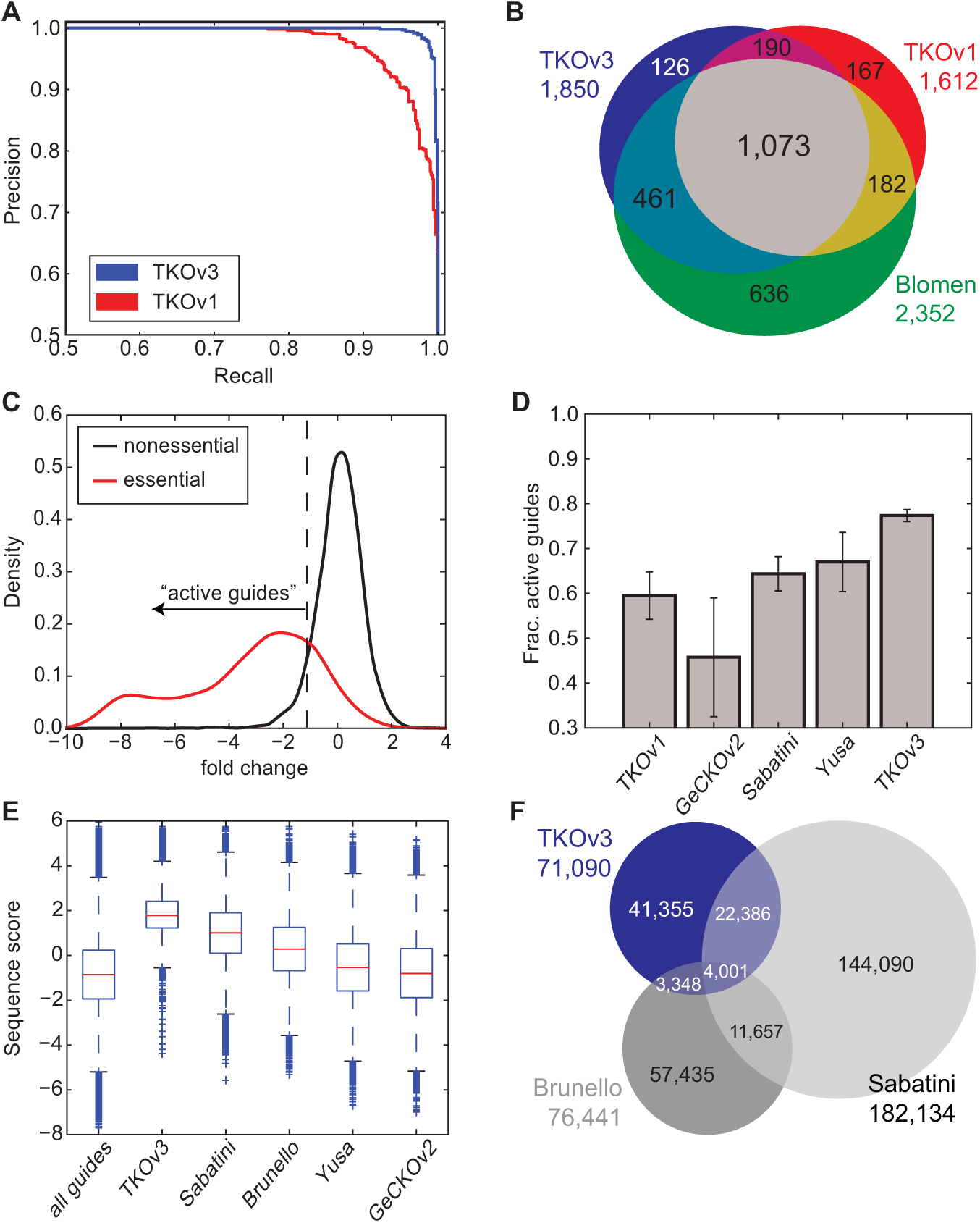
Evaluation of TKOv3 library. (A) Precision-recall curves of TKOv1 and TKOv3 screens in HAP1 cells. (B) Comparison of essential genes in TKOv3 vs. TKOv1 and HAP1 essentials from Blomen *et al.* at 5% FDR. (C) TKOv3 guides targeting essential vs. nonessential genes in HAP1. Guides targeting essential genes, with fold-change < 5^th^ percentile of guides targeting nonessential genes, are defined as ‘active guides’. (D) The fraction of active guides (active guides targeting essential genes / all guides targeting essential genes) across five libraries tested. (E) Distribution of sequence scores for all candidate gRNA sequences (n~2.5 million) compared to published CRISPR/Cas9 libraries. (F) Overlap of gRNA sequences in the top three libraries by sequence score.

As a further means of comparing library quality, we examined the observed dropout of guides targeting essential and nonessential genes. As expected, guides targeting reference nonessential genes showed a largely symmetric distribution of (log) fold-changes centered at zero (Fig 6c). We defined a scoring metric, the fraction of active guides, as the percentage of gRNAs targeting reference essential genes that show a fold-change greater than that of 95% of gRNAs targeting reference nonessential genes. The TKOv3 library shows a marked improvement over the TKOv1 library, as well as other latest-generation libraries for which screening data is available (Fig 6d). All of the libraries are substantially more efficient than GeCK0v2, as evaluated by data from [10]. This summary statistic is consistent with a precision-recall analysis of each library, performed by analyzing each screen with the BAGEL pipeline and using the same v2 reference essential (ie. CEG2) and reference nonessential gene sets.

Another library targeting human genes and designed from empirical observations is the Brunello library, described in [17]. We currently have no negative selection screen data from this library to evaluate the fraction of active guides, so as a proxy we calculated the distribution of guide sequence scores for all gRNAs in the library. Compared to all candidate gRNAs (all potential guide sequences targeting protein coding exons with an NGG PAM and 40-75% GC content; approx. 2.5 million sequences), the Brunello library has a higher sequence score than average, though not as high as either the Sabatini or TKOv3 libraries (Fig. 6e). However, it should be noted that low sequence score does not necessarily imply poor overall library performance; the Yusa library has sequence scores comparable to those in the GeCK0v2 library but due to its other design considerations (e.g. 3’ sequence bias, use of modified tracrRNA), it shows markedly better performance than GeCKOv2.

Given the similar sequence signatures of both TKOv3 and the Sabatini library (Fig. 6e), we examined the overlap of actual gRNA sequences between the two collections. Over 26,000 of the ~71,000 gRNA in TKOv3 are also in the Sabatini library, comprising some 37% of TKOv3 sequences. Both TKOv3 and Sabatini show markedly less overlap with the Brunello library (Fig 7e).

## Discussion

Gene knockout screening in mammalian cells is transforming human functional genomics and target discovery efforts, but CRISPR technology continues to evolve rapidly. Early proof-of-concept screens using pooled library approaches used large numbers of guide RNAs per gene to overcome the unknown sources of variation in gRNA targeting efficiency. We analyzed panels of pooled library screens from three different research groups, each using different CRISPR libraries, to identify a set of common hits across all tested conditions. This set of 684 genes, the “Core Essentials-2.0 or CEG2,” are consistent across adherent and suspension cell lines and represent a broader cross-section of essential cellular processes than the Core Essentials-1.0 genes derived from a panel of pooled library shRNA knockdown screens. Identification of CEG2 genes will be a useful metric for evaluating the sensitivity of genome-scale knockout screens in human cell lines.

Since the collection of known essential genes offers a set of expected outcomes for screens, we leveraged this knowledge to determine the characteristics of guide RNAs that maximize the discrimination of essential genes from nonessentials. We derived a sequence signature from the TKOv1 screens that predicts improved gRNA performance in TKOv1 screens as well as those using the GeCKOv2 and Sabatini libraries, though not the Yusa library. We also evaluated the effect of varying the number of gRNA per gene in the library, and observed that increasing library size beyond four gRNAs per gene typically yielded a small incremental increase in the sensitivity of the screen. Based on these observations we designed a new library, TKOv3, which contains four sequence-optimized guides targeting each of 18,053 protein-coding genes. The result is a library of 71,090 gRNA sequences that is small enough to facilitate genome-scale screens in cell lines while sensitive enough to minimize false negatives in a well-designed screen. TKOv3 library is a one-component library, expressing Cas9 from the viral vector, which relaxes the requirement to knock Cas9 into cell lines; however, we do not have a direct comparison of one-component and two-component libraries using the same optimized sequences.

Overall, improving the accuracy and scalability of CRISPR screens offers considerable benefits for the systematic survey of context-dependent essential genes across tissue types, genetic mutational landscapes, and environmental stimuli. Further efficiencies may be gained by exploring alternative Cas proteins, or engineering existing ones, for a variety of functions: to increase nuclease effectiveness, to broaden the addressable set of cleavage sites by using alternative PAM sequences, and especially by exploiting endogenous or engineered multiplexing capabilities. Though each alternative CRISPR-associated nuclease will almost certainly require a sequence optimization survey such as this one, the consistency of the latest-generation Cas9 libraries suggests that we are approaching a maximally efficient Cas9 gRNA design.

## Methods

The supplementary file HART_data_and_python_notebooks.tgz will be included as a hyperlink for download and contains python notebooks and all required data to generate the figures presented here. As such, it contains a near-complete, granular description of the computational methods applied in this study. Detailed experimental methods, and a summary of computational methods, are included below.

### Analysis of screens from various libraries

Guide RNA readcount data were downloaded from [2, 3, 5, 8]. Fold-changes were calculated by normalizing each sample to 10 million reads and calculating log2(experimental/control) for each replicate of each sample at each timepoint. Control was either the gRNA counts from the genomic DNA collected after infection (“TO”) or library plasmid pool, depending on the experimental design. A pseudocount of 0.5 reads was added to all readcounts to prevent discontinuities from zeros. gRNA with <30 reads at the T0 timepoint were excluded from the fold-change calculation.

Fold-changes were processed with BAGEL [14] using essential and nonessential training sets defined in [15]. The resulting Bayes Factors for all screens are included in Supplementary Table 1.

After the CEG2 set was defined, BFs for all screens were recalculated using this new reference set (Supplementary Table 2).

### Identification of core v2

Of the seventeen gRNA screens initially evaluated, three were withheld for later evaluation and analysis. Two others were excluded for relatively poor performance. For the remaining 12 screens, the Bayes Factor and the number of gRNAs targeting the gene were considered. Note that the number of gRNAs may vary by cell line and by library since only gRNAs with >30 reads in the T0 control sample were used for each cell line screen.

A gene was defined as “effectively assayed” if it was targeted by at least three gRNAs in a given screen. The CEG2 set was defined as genes effectively assayed in at least seven cell lines, with a Bayes Factor >=6 in 85% of cell lines in which they were assayed. Since most genes were assayed in either 7 or 12 cell lines, this effectively means that core essentials are hits in 6 of 7 or 11 of 12 screens.

### Evaluation of gRNA/gene

The Sabatini library in Wang et al [5] contains 10 gRNA per gene. For each of the five screens in four cell lines (KBM7 is screened twice), a subset of guides were randomly selected and Bayes Factors were calculated from the subset. This process was repeated 10 times for each count of n=2 to n=7 gRNA per gene. Performance was evaluated by counting the fraction of core essentials recovered and the overall number of hits called at a defined threshold.

### Evaluation of replicates/screen

The TKOv1 screen in RPE1 cells[2] and Yusa library screen in HL60 cells [8] were conducted with similar three-replicate experimental designs. We ran BAGEL on each replicate independently, and on all three combinations of two replicates, and evaluated performance of each as in gRNA/gene above.

### Nontargeting vs. nonessential controls

The Sabatini library contains ~1,000 nontargeting gRNA controls [5]. We compared the distribution of fold-changes for gRNA nontargeting controls to the distribution of fold-changes for gRNA targeting gold standard nonessential genes. Statistical significance was calculated by T-test.

### Identifying sequence signature

To identify a sequence signature that predicts high-performing guides, we evaluated data from TKOv1 screens. From the base 90k TKOv1 library [2], we identified genes in the new Core Essentials-2.0 set that were targeted by six gRNAs each. gRNAs were ranked by log fold-change, and the three gRNAs with the “best” (most negative) fold-change were identified, as well as the “worst” (remaining three gRNA). Then, the frequency of each nucleotide at each position in the 20mer guide sequence was calculated for all “best” guides targeting all selected genes, and the same was done for the “worst” guides. The “worst” frequency was subtracted from the “best,” resulting in a delta-frequency table.This process was repeated independently for each replicate at the endpoint for six TKOv1 screens (DLD1, GBM, HAP1, HCT116, RPE1, RPE1dTP53) for a total of 16 samples.

The delta-frequency tables were summed across the 16 samples and scaled so that the most extreme value (C18) equals one. As TKOv1 explicitly excludes gRNA with T in the last four positions, no score is discovered here; we manually set the score to - 1 at these four positions. The final score table is in Supplementary Table 4. To calculate the sequence score of any candidate gRNA sequence, simply add the nucleotide scores at each position of the gRNA.

The score table was evaluated against the 85k supplementary TKOv1 library. We calculated the sequence score for all gRNA targeting essential genes, then compared the fold-change distribution of gRNA in the top quartile of scores to the gRNA in the bottom quartile. We repeated this process for the Yusa, Sabatini, and GeCKO v2 libraries.

### TKOv3 library design

Gene models for protein-coding genes were derived from Gencode v19 gene models (all genomic analysis was done with hg19). Coding exons were numbered in ascending order from the transcription start site.

The genome was scanned and a list of all candidate gRNA targeting all genes was generated, for all candidate gRNA such that the Cas9 cut site would be in a coding region. Candidate gRNAs were then filtered for the following criteria: 40-75% GC content, no homopolymers of length of 4 or greater, no restriction sites for AgeI (ACCGGT), KpnI (GGTACC), BveI/BspMI (ACCTGC), BsmI (GAATGC), or BsmBI (CGTCTC), and no common SNP (db138) in either the PAM or the protospacer sequence.

Remaining gRNAs were mapped to hg19 with Bowtie (with NGG PAM sequence appended), allowing up to two mismatches outside the PAM “n” position. Guide sequences were excluded if there was any match within either exonic or intronic sequences of an off-target gene. Guide sequences were then assigned to class 1 in the very rare case where the sequence matches the target gene more than once and does not have any predicted off-target cut sites. Remaining guides were assigned as class 2 (hits targeted gene once; no off-targets; common), class 3 (up to two off-target hits within two mismatches, if off-target sites are in intergenic regions; very common), or class 4 (up to 3 off-target intergenic hits). The sequence score was also calculated for each guide, with the median sequence score across all guides being ~ - 1. In general, a small number of intergenic off-target cut sites has been shown to have negligible impact on cellular fitness relative to guides targeting known nonessential genes [2, 3]. The TKOv3 library exploits this observation by allowing one to two predicted intergenic off-target cut sites if the sequence score of the primary target site is particularly high.

Guides were further binned into ranks:

| **<u>Rank</u>** | **<u>Class</u>** | **<u>SeqScore</u>** | **<u># Candidate Guides</u>** |
| --- | --- | --- | --- |
| 1 | 1 | >0.0 | 1,501 |
| 2 | 1,2,3 | >0.85 | 286,415 |
| 3 | 1,2,3,4 | >0.85 | 307,059 |
| 4 | 1,2,3 | 0.0-0.85 | 304,477 |
| 5 | 1,2,3,4 | -1.0-0.85 | 837,136 |

To select guides for the library, we began with the rank1 candidate gRNAs. Then for each exon, we selected the single candidate gRNA with the top sequence score and added it to the library. This process was repeated four times, to allow up to four gRNAs per gene, with the competing goals of maximizing gRNA quality and exon coverage while allowing multiple gRNAs per exon if the gRNAs are high quality. The process therefore selects high-scoring guide sequences for four different exons, if available. If not, high-scoring guides targeting already-targeted exons are prioritized above low-scoring guides targeting different exons. These steps were repeated for each subsequent rank, with the following results

| **<u>Rank</u>** | **<u>Num. gRNA added</u>** | **<u>Cumulative gRNA in library</u>** |
| --- | --- | --- |
| 1 | 679 | 679 |
| 2 | 63,834 | 64,513 |
| 3 | 1,069 | 65,582 |
| 4 | 3,476 | 69,058 |
| 5 | 1,890 | 70,948 |

An additional 142 control sequences targeting EGFP, LacZ, and luciferase were also added, for a final library size of 71,090 gRNA.

### Genome-scale lentiviral gRNA construction

All 72,090 gRNAs were synthesized as 58mer oligonucleotides on one microarray chip (Custom Array) and amplified by PCR as a pool. The PCR products were purified using QIAquick nucleotide removal kit (Qiagen) and cloned into a modified version of the all-in-one lentiviral vector lentiCRISPRv2 (Addgene) which contains the spCas9 gene. The lentiCRISPRv2 vector was digested with BsmBI (NEB) for 2 hours at 55°C, treated with alkaline phosphatase (NEB) for 30 minutes at 37 °C and gel purified using the QIAquick Gel Extraction kit (Qiagen). Using a one-step digestion and ligation reaction, 40 fmoles of the purified library PCR pool was cloned into 7.1 fmoles of the digested lentiCRIPSRv2 vector for a ratio of 1:6 vector to insert molar ratio. The ligation reaction was purified using the QIAquick nucleotide removal kit (Qiagen) and 4 μl of the purified ligation was transformed into One Shot Stbl3 competent cells (Thermo Fisher Scientific). To yield a x-fold representation of the library, 40 identical ligation reactions were pooled and purified followed by 40 parallel transformations. Outgrowth media from transformations were pooled and plated onto 100 15cm LB-carbenicillin (100 μg/ml) plates. Colonies were scraped off plates, pooled and the plasmid DNA was extracted using the QIAfilter Plasmid Giga kit (Qiagen).

### Cell culture

HEK293T cells were maintained in Dulbecco’s modified Eagle’s medium (DMEM) with high glucose and pyruvate (Thermo Fisher Scientific) supplemented with 10% FBS (Thermo Fisher Scientific) and 1% Penicillin/Streptomycin (Thermo Fisher Scientific). HAP1 cells were obtained from Horizon Discovery and maintained in Iscove’s Modified Dulbecco’s Medium (IMDM) supplemented with 10% FBS and 1% Penicillin/Streptomycin. All cells were maintained in humidified incubators at 37°C and 5% CO_2_.

### Lentivirus production

TKOv3 library lentivirus was produced by co-transfection of lentiviral vectors psPAX (packaging vector) and pMDG.2 (envelope vector) with TKOv3 lentiCRISPR plasmid library using X-tremeGene™ 9 transfection reagent (Roche). Briefly, HEK293T cells were seeded at a density of 9×l0^6^ cells per 15 cm plate and incubated overnight, after which cells were transfected with a mixture of psPAX (4.8μg), pMDG.2 (3.2μg), TKOv3 plasmid library (8μg) and X-tremeGene™ 9 (48μl) in accordance with the manufacturer’s protocol. 24 hours post transfection the medium was changed to serum-free, high BSA growth medium (DMEM, 1% BSA,1% Penicillin/Streptomycin). Virus containing medium was harvested 48 hours after transfection, centrifuged at 1,500 rpm for 5 minutes and stored at −80°C. Functional titers in HAP1 cells were determined by infecting cells with a titration of TKOv3 lentiviral library in the presence of polybrene (8μg/ml). 24 hours after infection medium was replaced with puromycin (2 μg/ml) containing medium to select for transduced cells and incubated for 48 hours. The multiplicity of infection (MOI) of the titrated virus was determined 72-hours post infection by comparing the percent survival of infected cells to non-infected control cells.

### Pooled genome-wide CRISPR drop out screens in HAP1 cells

50×l0^6^ HAP1 cells were infected with TKOv3 lentiviral library (71,090 gRNAs) at a MOI of ~0.3 to achieve approximately 200-fold coverage of the library after selection. 72 hours after infection, selected cells were split into three replicates containing 15×l0^6^ cells each, passaged every 3-4 days and maintained at 200-fold coverage. 15×l0^6^ cells were collected for genomic DNA extraction at day 0 and at every passage until day 18 post selection, or approximately 15 doublings.

Genomic DNA was extracted from cell pellets using the QIAamp Blood Maxi Kit (Qiagen), precipitated using ethanol and sodium chloride and resuspended in EB buffer. gRNA inserts were amplified via PCR using primers harboring Illumina TruSeq adapaters i5 and i7 barcodes and the resulting libraries were sequenced on an Illumina HiSeq2500. Each read was completed with standard primers for dual indexing with Rapid Run V1 reagents. The first 20 cycles of sequencing were “dark cycles”, or base additions without imaging. The actual 26bp read begins after the dark cycles and contains two index reads, reading the i7 first, followed by i5 sequences.

## Acknowledgments

We wish to thank members of the Moffat, Hart, Durocher, and Angers labs for helpful discussions. TH was supported by MD Anderson Cancer Center Support Grant P30 CA016672 (the Bioinformatics Shared Resource) and the Cancer Prevention Research Institute of Texas (CPRIT) grant RR160032. NH is a long-term fellow of the Human Science Frontier Program. KL holds a Canada Vanier Graduate Scholarship, MA holds a postdoctoral fellowship award from the Swiss National Science Foundation, and MC holds an Ontario Graduate Scholarship. This work was supported from grants funded by the Canadian Institutes for Health Research to DD (#703893) and JM (MOP-142375), Canadian Cancer Society Research Institute to DD (#703893), and the Ontario Research Fund to BA, CB and JM (ORF-RE7-037). DD and CB are a Canada Research Chairs (Tier 1) in the Molecular Mechanisms ofGenome Integrity and Genetics, respectively. JM is a Canada Research Chair (Tier 2) in Functional Genetics.

## References

1. Jinek, M., et al.,A programmable dual-RNA-guided DNA endonuclease in adaptive bacterial immunity. Science, 2012. 337(6096): p. 816–21.

2. Hart, T., et al.,High-Resolution CRISPR Screens Reveal Fitness Genes and Genotype-Specific Cancer Liabilities. Cell, 2015.163(6): p. 1515–26.

3. Koike-Yusa, H., et al.,Genome-wide recessive genetic screening in mammalian cells with a lentiviral CRISPR-guide RNA library. Nature biotechnology, 2014. 32(3): p.267–73.

4. Shalem, O., et al.,Genome-Scale CRISPR-Cas9 Knockout Screening in Human Cells. Science, 2013.

5. Wang, T., et al.,Identification and characterization of essential genes in the human genome. Science, 2015.

6. Wang, T., et al.,Genetic Screens in Human Cells Using the CRISPR/Cas9 System. Science, 2013.

7. ParnasO, J.M., Eisenhaure TM, Herbst RH,, et al.,A genome-wide CRISPR screen in primary immune cells to dissect regulatory networks. Cell, 2015.

8. Tzelepis, K., et al.,A CRISPR Dropout Screen Identifies Genetic Vulnerabilities and Therapeutic Targets in Acute Myeloid Leukemia. Cell Rep, 2016.17(4): p. 1193–1205.

9. Wang, T., et al.,Gene Essentiality Profiling Reveals Gene Networks and Synthetic Lethal Interactions with Oncogenic Ras. Cell, 2017.

10. Aguirre, A.J., et al.,Genomic Copy Number Dictates a Gene-Independent Cell Response to CRISPR/Cas9 Targeting. Cancer Discov, 2016. 6(8): p. 914–29.

11. Evers, B., et al.,CRISPR knockout screening outperforms shRNA and CRISPRi in identifying essential genes. Nat Biotechnol, 2016. 34(6): p. 631–3.

12. Kaelin, W.G., Jr., Molecular biology. Use and abuse of RNAi to study mammalian gene function. Science, 2012. 337(6093): p. 421–2.

13. Echeverri, C.J., et al.,Minimizing the risk of reporting false positives in large-scale RNAiscreens. Nat Methods, 2006. 3(10): p. 777–9.

14. Hart, T. and J. Moffat, BAGEL: a computational framework for identifying essential genes from pooled library screens. BMC Bioinformatics, 2016.17: p. 164.

15. Hart, T., et al.,Measuring error rates in genomic perturbation screens: gold standards for human functional genomics. Molecular systems biology, 2014. 10: p.733.

16. Steinhart, Z., et al.,Genome-wide CRISPR screens reveal a Wnt-FZD5 signaling circuit as a druggable vulnerability of RNF43-mutant pancreatic tumors. Nat Med, 2017. 23(1): p.60–68.

17. Doench, J.G., et al.,Optimized sgRNA design to maximize activity and minimize off-target effects of CRISPR-Cas9. Nat Biotechnol, 2016. 34(2): p. 184–91.

18. Haeussler, M., et al.,Evaluation of off-target and on-target scoring algorithms and integration into the guide RNA selection tool CRISPOR. Genome Biol, 2016. 17(1): p. 148.

19. Heigwer, F., G. Kerr, and M. Boutros,E-CRISP: fast CRISPR target site identification. Nat Methods, 2014.11(2): p. 122–3.

20. Heigwer, F., et al.,CRISPR library designer (CLD): software for multispecies design of single guide RNA libraries. Genome Biol, 2016.17: p. 55.

21. Hsu, P.D., et al.,DNA targeting specificity ofRNA-guided Cas9 nucleases. Nat Biotechnol, 2013. 31(9): p. 827–32.

22. Liu, X., et al.,Sequence features associated with the cleavage efficiency of CRISPR/Cas9 system. Sci Rep, 2016. 6: p. 19675.

23. Park, J., J.S. Kim, and S. Bae, Cas-Database: web-based genome-wide guide RNA library design for gene knockout screens using CRISPR-Cas9. Bioinformatics, 2016. 32(13): p.2017–23.

24. Blomen, V.A., et al.,Gene essentiality and synthetic lethality in haploid human cells. Science, 2015. 350(6264): p. 1092–6.

